# Astrocytes improve neuronal health after cisplatin treatment through mitochondrial transfer

**DOI:** 10.1101/2019.12.17.879924

**Authors:** Krystal English, Andrew Shepherd, Ndidi-Ese Nzor, Ronnie Trinh, Annemieke Kavelaars, Cobi J. Heijnen

**Affiliations:** Laboratories of Neuroimmunology, Department of Symptom Research, Division of Internal Medicine, The University of Texas MD Anderson Cancer Center, Houston, TX 77030; Department of Neurobiology & Anatomy, The University of Texas McGovern Medical School, Houston, TX 77030; Department of Neurology, The University of Texas McGovern Medical School, Houston, Tx 77030

## Abstract

Neurodegenerative disorders, including chemotherapy-induced cognitive impairment, are associated with neuronal mitochondrial dysfunction. Cisplatin, a commonly used chemotherapeutic, induces neuronal mitochondrial dysfunction *in vivo* and *in vitro*. Astrocytes are key players in supporting neuronal development, synaptogenesis, axonal growth, metabolism and, potentially, mitochondrial health. We tested the hypothesis that astrocytes transfer healthy mitochondria to neurons after cisplatin treatment to restore neuronal health. We used an *in vitro* system in which astrocytes containing mito-mCherry-labeled mitochondria were co-cultured with primary cortical neurons damaged by cisplatin. Culture of primary cortical neurons with cisplatin reduced neuronal survival and depolarized neuronal mitochondrial membrane potential. Cisplatin induced abnormalities in neuronal calcium dynamics characterized by increased resting calcium levels, reduced calcium responses to stimulation with KCl, and slower calcium clearance. The same dose of cisplatin that caused neuronal damage did not affect astrocyte survival or astrocytic mitochondrial respiration. Co-culture of cisplatin-treated neurons with astrocytes increased neuronal survival, restored neuronal mitochondrial membrane potential, and normalized neuronal calcium dynamics especially in those neurons that had received mitochondria from astrocytes. These beneficial effects of astrocytes were associated with transfer of mitochondria from astrocytes to cisplatin-treated neurons. The Rho-GTPase Miro-1 is known to contribute to mitochondrial motility and transfer. We show that siRNA-mediated knockdown of Miro-1 in astrocytes reduced mitochondrial transfer from astrocytes to neurons and prevented the normalization of neuronal calcium dynamics.

In conclusion, we identified transfer of mitochondria from astrocytes to neurons damaged by cisplatin as an important repair mechanism to protect cortical neurons against the toxic effects of this chemotherapeutic.

**Significance statement:** Chemotherapy-induced neurotoxicity is a serious health problem and little is known about the underlying mechanisms. Especially neurons are very sensitive to cisplatin treatment. We show that astrocytes can protect neurons damaged by cisplatin by improving neuronal survival, mitochondrial health, and calcium dynamics *in vitro*. This beneficial effect of astrocytes is dependent on the transfer of mitochondria from astrocytes to the damaged neurons. Our findings provide evidence for an important endogenous protective neuro-glial mechanism that could contribute to prevention of neuronal death as a result of cisplatin treatment and thereby aid in sustaining brain health of patients during chemotherapy.

## Introduction

Mitochondria are unique organelles that are crucial for sustaining cellular health through multiple functions, including ATP production via oxidative phosphorylation, metabolic regulation, regulation of apoptosis, and Ca^2+^ buffering (Kann and Kovács, 2007; Burté *et al*., 2015; Kaasik, 2015; Flippo and Strack, 2017). Neurons are highly specialized cells that, like all cells, are critically dependent on intact mitochondrial function for rapidly responding to changes in energy demand, storing and buffering Ca^2+^ and, specifically for neurons, neurotransmission and plasticity (McCue, Haynes and Burgoyne, 2010; Pivovarova and Andrews, 2010; Guo, Tian and Du, 2017; Smith and Gallo, 2018). Due to the critical importance of mitochondria for multiple key aspects of neuronal function, it is not surprising that mitochondrial dysfunction can have devastating effects on brain function (Chiu *et al*., 2017; Devine and Kittler, 2018; Ren *et al*., 2019). Astrocytes release multiple factors that are essential to neuronal development, signaling, metabolism, axonal growth and synaptogenesis (Bayraktar et al. 2014; Bazargani and Attwell 2016; Bozoyan, Khlghatyan, and Saghatelyan 2012; Gengatharan, Bammann, and Saghatelyan 2016; Lundgaard et al. 2014). Recent evidence indicates an additional way via which astrocytes can contribute to neuronal health is by donating healthy mitochondria to damaged neurons (Vignais *et al*. 2017; Wang *et al*. 2011). Specifically, Wang *et al*. (2011) showed that exposure of rat hippocampal astrocytes and neurons to H_2_O_2_ or serum deprivation promotes transfer of mitochondria from astrocytes to neurons. There is also evidence from *in vitro* and *in vivo* models of ischemic stroke that astrocytes release vesicles containing mitochondria that can be taken up by neurons (Berridge, Schneider and Mcconnell, 2016; Hayakawa *et al*., 2016). Moreover, astrocytes do not only function as donor of healthy mitochondria but can also function as recipient of damaged mitochondria. Davis *et al*. (2014), showed that retinal ganglion cell axons routinely shed mitochondria at the optic nerve head to be degraded by astrocytes *in vivo*. Mitochondrial transfer from one cell type to another is not exclusive to astrocytes and neurons. For example, we and others have shown that mesenchymal stem cells transfer mitochondria to damaged neuronal stem cells thereby improving stem cell survival and mitochondrial membrane potential in the recipient cells (Boukelmoune *et al*., 2018; Gheusi and Lledo, 2014; M. L. Vignais *et al*., 2017; Babenko *et al*., 2018). Transfer of mitochondria from one cell to another occurs via multiple mechanisms such as release and uptake of vesicles, transfer via gap junctions, and transfer via F-actin based tunneling nanotubes (Rogers and Bhattacharya, 2013; M.-L. Vignais *et al*., 2017). Mitochondrial Rho-GTPase 1 (Miro-1) is a calcium-sensitive adaptor protein that drives movement of mitochondria along microtubules (Jackson and Robinson, 2015). Miro-1 is involved in transferring mitochondria from mesenchymal stem cells to neuronal stem cells (Boukelmoune *et al*., 2018), but its contribution to mitochondrial transfer from astrocytes to neurons is unknown.

Multiple neurodegenerative disorders, including Parkinson’s disease, Alzheimer’s disease, and chemotherapy-induced cognitive impairment are associated with neuronal mitochondrial dysfunction (Devine and Kittler, 2018; Ma *et al*., 2018; Ren *et al*., 2019, Chiu *et al*., 2016). We have shown recently that treatment of mice with the chemotherapeutic drug cisplatin results in synaptosomal mitochondrial dysfunction that causes cognitive deficits (Chiu *et al*., 2016). Cisplatin crosses the blood-brain barrier at levels that are sufficient to cause damage to hippocampal neurons and to neuronal stem cells (Andres *et al*., 2014). However, cisplatin treatment *in vivo* does not lead to overt neuronal cell death which could indicate that there are endogenous protective mechanisms, such as mitochondrial transfer by astrocytes, to assist in sustaining neuronal health in conditions of acute danger to adult neurons.

The aim of the current study is to test the hypothesis that astrocytes transfer mitochondria to neurons damaged by cisplatin and thereby improve neuronal function and health *in vitro*.

## Methods

### Culture of cortical neuron and astrocytes

Timed-pregnant Long Evans rats (Charles River, Wilmington, MA, USA) were sacrificed and E18 fetuses of both sexes were collected in accordance with Institutional Animal Care and Use Committee-approved protocols. Cortices were dissected and incubated in 10 ml of dissociation media (81.8mM Na_2_So_4_, 30mM K_2_SO_4_, 5.8mM MgCl_2_, 0.25mM CaCl_2_, 1mM Hepes, 20mM glucose, 0.0001% Phenol Red, 0.16mM NaOH pH=7.4) that contained 10 U/mL papain and 5mg of L-cysteine in total (Worthington, Lakewood, NJ, USA) for 10 minutes at 37°C, followed by incubation with 150mg of trypsin inhibitor (Millipore-Sigma, St. Louis, MO) in 10ml of dissociation media for 10 minutes at 37°C. The cortical tissue was mechanically dissociation in Opti-mem (GIBCO, Carlsbad, CA, USA) with 2.5M Glucose (GIBCO). Cells were cultured on plates coated with 0.05mg/ml poly-D-lysine (PDL; Millipore-Sigma) in neurobasal medium (NBM) with 100 U/mL penicillin and 1x B-27 supplement (Invitrogen Carlsbad, CA) at 37°C and 5% CO_2_. Neuronal cultures were maintained in NBM with B-27 supplement and media was replaced every 3 days. Cortical astrocytes were grown in DMEM/F12 medium supplemented with 10% fetal bovine serum and 5% of 10,000 units/ ml of penicillin and 10,0000 µg/ml of streptomycin (GIBCO) at 5% CO_2_ and 37 °C.

Neuronal cells were used for experiments after 12-15 days of culturing *in vitro* (DIV). To confirm neuronal enrichment, cells were fixed with 4% paraformaldehyde in PBS, treated with 0.25% Triton X-100, blocked in 2% BSA in PBS and stained with anti-Map2 antibody (1:2000); Sigma-Aldrich); anti-GFAP (1:200, Acris, Rockville, MD) and anti-Olig2 (1:400, Abcam, Cambridge, UK) antibody. On DIV12 > 98% of cells were Map2+ and Olig2 and GFAP-negative. Astrocytes were used until the third passage and were >99% GFAP+.

### Astrocyte transfections

Astrocytes were plated in a 6-well plate at 1.5×10^5^ cells/well 24 hours before transfection. For labeling mitochondria, astrocytes were transfected with 2 μg of pLYS1-FLAG-MitoGFP-HA (Addgene plasmid # 50057) which contains the pore-forming subunit of the mitochondrial calcium uniporter coupled to GFP or a mito-mCherry construct generated by subcloning the targeting sequence of the pLYS1-FLAG-MitoGFP-HA plasmid into the mcherry2-N1 vector (Addgene plasmid # 54517). For Miro-1 knockdown, 5nmol of Rho-1 siRNA (Qiagen, Germany #SI01401743) was diluted in rnase-free water (provided in kit) to make a 20uM stock, andAllStars Negative Control Scrambled siRNA (Qiagen #SI03650318) was performed similarly to make a 20uM stock instructions provided. Astrocytes were transfected with 80 nM of Rho-1 siRNA from a 20uM stock (Qiagen, Germany #SI01401743) or 80 nM AllStars Negative Control Scrambled siRNA from a 20uM stock (Qiagen #SI03650318). All transfections were performed using the Astrocyte Transfection kit (Altogen Biosystems, Las Vegas, NV, USA) according to manufacturer’s instructions. Miro-1 knockdown was confirmed by Western blot with anti-Rho1 antibody (Novus Biologicals, Centennial, CO, USA) with GAPDH as control (Abcam) followed detection of bands with enhanced chemoluminescence (GE Healthcare Bio-Sciences, Pittsburgh, PA). Blots were captured in the LAS system using Image Quant software (GE Healthcare Bio-Sciences) for quantification of bands.

### Analysis of neuronal survival

Neurons were plated in 96 well plates coated with 0.05mg/ml PDL at 5×10^4^ neurons/well. Viability after exposure to cisplatin (Teva, Petah Tikva, Israel) was quantified using the colorimetric cell viability reagent WST-1 (Millipore-Sigma, #11644807001). To assess the effect of astrocytes on neuronal survival, separate cultures of neurons (1.5 × 10^5^ cells/well in a 6-well plate) and astrocytes (5×10^4^ /well in a 6-well plate) were treated with cisplatin for 24 h. Neurons were then labeled with 20 μM CellTracker Blue (CTB; Invitrogen) for 45 min at 37 °C, and washed in serum-free media. Astrocytes (5×10^4^ cells/well) were added to the neuronal culture and survival of CTB+ neurons was quantified 17 hours later using a Countess II FL automated cell counter (Invitrogen).

### Analysis of mitochondrial membrane potential and mitochondrial transfer

Neurons were plated at 1.5×10^5^ cells/well in a 6-well plate, treated with cisplatin for 24 h, and labeled with 20µM CellTracker Green (Thermo Fisher). Astrocytes (5×10^4^cells/well) were added and co-cultured with the neurons for 17 h. The co-cultures were stained with tetramethylrhodamine methyl ester (TMRM, Invitrogen; 250 nM) for 45 min at 37 °C, or Mitotracker (50nM, Thermo Fisher, Waltham, MA, USA) for 30 min at 37 °C. As a positive control, neurons were treated with 10 μM carbonilcyanide p-triflouromethoxyphenylhydrazone (FCCP, Sigma-Aldrich), a mitochondrial uncoupler, for 15 min. Cells were collected and TMRM fluorescence intensity of the cell tracker green positive cells was quantified using an Accuri C6 Flow Cytometer (BD Biosciences, San Jose, CA USA). For confocal microscopy, neurons were plated in cell culture imaging dishes (ibidi, Fitchburg, WI, USA), treated with cisplatin, stained with CTB and cultured with or without mito-GFP- and cell tracker deep red-labeled astrocytes for 17 h followed by staining with TMRM or Mitotracker. The TMRM was used in sub-quench mode as described by Rego et al., (2001).

For analysis of mitochondrial transfer, neurons exposed to cisplatin or vehicle were labeled with 20µM CTB (Thermo Fisher) prior to co-culture with astrocytes. Astrocytes were transfected with either mito-GFP or mito-mCherry prior to exposure to cisplatin followed by labeling with 20µM Celltracker DeepRed (Thermo Fisher) and co-culture with neurons. The co-cultures were imaged on an SPE Leica Confocal Microscope (Leica Microsystems, Buffalo Grove, IL, USA) with a 63 X or 40 X objective and images were analyzed with LAS X software.

### Analysis of mitochondrial bioenergetics

To assess mitochondrial bioenergetics, astrocytes (5×10^4^ cells/well) were plated in a Seahorse XFe 24 microplate (Seahorse Biosciences/Agilent Technologies, Santa Clara, CA, USA) coated with 0.05mg/ml PDL and treated with 1 μM cisplatin or vehicle for 24h. Cells were washed and incubated for 1 h at 37 °C in XF base media (Seahorse Biosciences) supplemented with 11 mM glucose (Sigma-Aldrich), 2 mM glutamine (Sigma Aldrich), and 1 mM pyruvate (Sigma-Aldrich), 2 mM Oligomycin (Sigma-Alrich), 4 mM FCCP, and rotenone/antimycin A (Sigma-Aldrich, 2 mM each) were used with a 3-time repeat of a 2-min mix, 3-min wait, and 2-min measure assay cycle. Oxygen consumption rates were normalized to the total protein content of each well. Basal respiration, maximal respiratory capacity, and spare respiratory capacity were determined as described previously (Boukelmoune *et al*., 2018).

### Calcium imaging

Functional Ca^2+^ imaging on cortical neurons was performed as described previously (Shepherd *et al*., 2018). 12mm circular glass coverslips containing cells were incubated at room temperature for 20 min with the Ca^2+^-sensitive dye Fura-2-AM (Invitrogen, 2 μM), dissolved in standard extracellular HEPES-buffered HBSS (known hereafter as extracellular imaging buffer) containing the following (in mM): 140 NaCl, 5 KCl, 1.3 CaCl_2_, 0.4 MgSO_4_, 0.5 MgCl_2_, 0.4 KH_2_PO_4_, 0.6 NaHPO_4_, 3 NaHCO_3_, 10 glucose, and 10 HEPES adjusted to pH 7.4 with NaOH and 310 mOsm with sucrose. The coverslip was placed in the recording chamber (ALA scientific Instruments, Farmingdale, NY, USA) mounted on the stage of an inverted Nikon Ti2 microscope and continuously superfused for 5 min at room temperature with extracellular imaging buffer. Fura-2 fluorescence was alternately excited at 340 and 380 nm (12 nm band-pass, 50ms exposure) at 1Hz using a Lambda LS Xenon lamp (Sutter Instruments, Novato, CA, USA) and a 10x/NA 0.5 objective or 40x/NA 0.6. The CFI Super Fluor 10X was used to measure the ratio of Fura-2 fluorescence.Emitted fluorescence was collected at 510 nm using a sCMOS pco.edge camera for the entire experimental duration, including the first 5 min wash duration, and the ratio of fluorescence (F340:F380) was calculated. The shift in the ratio of Fura-2 fluorescence between excitation at 340 nm versus 380 nm is used as a readout of changes in intracellular calcium concentration [Ca^2+^_i_]. Baseline recording of F340:F380 ratio without stimulus was completed to obtain an indication of resting [Ca^2+^_i_]. Neurons were stimulated with 20 mM of KCl in the extracellular imaging buffer with continuous superfusion at room temperature. Imaging was analyzed with the NIS Elements software. To identify neurons that did or did not receive astrocytic mitochondria, the Nikon inverted Ti2 microscope with associated A1Rsi-HD confocal system and NIS Elements software was used with the CFI S Plan Fluor ELWD 40XC objective. A single exposure at 576nm excitation was used to quantify fluorescence within neurons before beginning measurement of F340:F380 ratio.

### Data analysis

Data are presented as mean ± SEM of at least 3 independent experiments performed in duplicate or triplicate. For survival analysis and mitochondrial membrane potential analysis, data were normalized to the control of each experiment and replicates were averaged. We used One-way or Two-way analysis of variance (ANOVA) with or without repeated measure followed by Tukey’s correction for multiple comparisons or Sidak’s correction for multiple comparisons according to experimental set up. For mitochondrial transfer analysis, the percentage of neurons that received mitochondria was calculate for each group. We used Student’s t-test or Two-way analysis of variance (ANOV) without repeated measures followed by Tukey’s correction for multiple comparisons. For calcium imaging, the ratio of fluorescence (F340:F380) was calculated and the ratios at baseline were subtracted from the maximum peak values upon KCl stimulus to calculate change in [Ca^2+^_i_]. To calculate Ca^2+^ clearance, F340:F380 ratios were converted to a 0-1 scale with 1 being the maximum value in response to 20mM KCl. Then, we quantified the time to return to 20% of baseline by counting the values from 1 (maximum peak) to 20% of the baseline value for each neurons. All analyses were performed using GraphPad Prism 8 (GraphPad Software, La Jolla, CA, USA).

## Results

### Astrocytes improve neuronal survival after cisplatin treatment

To examine the effect of cisplatin on neuron and astrocyte survival, we treated separate cultures of primary cortical neurons and astrocytes with cisplatin for 24 h and measured survival using the WST assay. Cisplatin dose-dependently reduced neuronal survival as assessed immediately after removal of cisplatin (Figure 1A) or 17 hours later (Figure 1B). Astrocytes were shown to be resistant to cisplatin since only the highest dose (4 μM) resulted in a modest reduction in astrocyte survival (Figure 1C and D). For all further experiments we used cisplatin at a dose of 1 μM. Exposure of astrocytes to 1 μM cisplatin did not induce detectable changes in astrocytic mitochondrial respiration; basal respiration, spare respiratory capacity and maximal respiration were similar in control and cisplatin-treated astrocytes (Figure 1E). To test the hypothesis that astrocytes improve the survival of neurons treated with cisplatin, primary cortical neurons and astrocytes were cultured separately in the presence or absence of 1 μM cisplatin for 24 h. Next cisplatin was removed, neurons were labeled with celltracker blue (CTB), and co-cultured with astrocytes for an additional 17 h. Co-culture with astrocytes significantly improved survival of cisplatin-treated neurons. Astrocytes did not affect survival of control neurons (Figure 1F).

**Figure 1:**
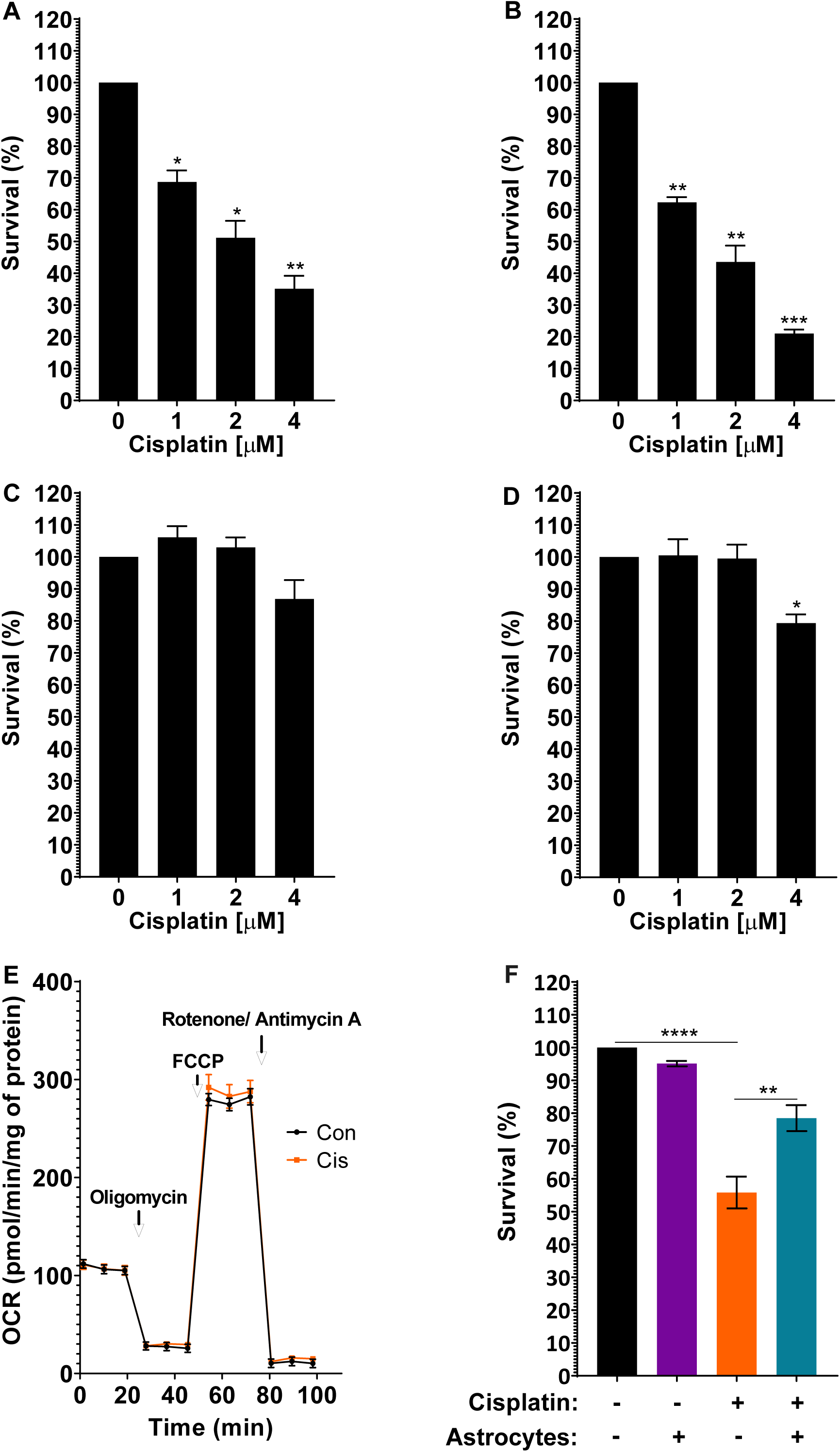
Astrocytes improve survival of neurons damaged by cisplatin. A-D. Survival of neurons (A, B) and astrocytes (C, D) after exposure to cisplatin. Primary cultures of cortical neurons (12 days *in vitro*) or cortical astrocytes were treated with increasing doses of cisplatin or vehicle for 24 h, then the cisplatin was removed and replaced with media. Survival was measured using a WST-1 assay: (A, C) at the end of the 24 h culture period or (B, D) 17 h after removal of cisplatin. Data was normalized to survival in the absence of cisplatin and analyzed using one-way ANOVA followed by Dunnett’s multiple comparisons test (\**p*<*0*.*05*, \*\**p*<*0*.*01*, \*\*\**p*<*0*.*001*). E. Astrocytes were treated with 1µM cisplatin for 24 h followed by 17 h of culture without cisplatin. Oxygen consumption rates (OCR) were analyzed using Seahorse XFe 24 Analyzer and normalized to protein content. F. Neurons and astrocytes were treated with cisplatin or control medium for 24 h. Cisplatin was removed, neurons were labeled with Celltracker Blue (CTB) and cultured for another 17 h with or without astrocytes, and the number of surviving neurons was counted. Data are normalized to control and represents the mean ± SEM of 3 independent experiments performed in duplicate. Two-way ANOVA: cisplatin x astrocyte interaction: \*\**p*<*0*.*01*; Tukey’s multiple comparisons post hoc test (\**p*<*0*.*05*, \*\**p*<*0*.*01*, \*\*\*\**p*<*0*.*0001*).

### Astrocytes improve mitochondrial membrane potential of neurons damaged by cisplatin

To assess the effect of cisplatin on neuronal mitochondrial integrity, we quantified mitochondrial membrane potential by labeling the cells with tetramethylrhodamine (TMRM) and assessed fluorescence intensity by flow cytometry. Figure 2A clearly shows that cisplatin depolarized the neuronal mitochondrial membrane potential as indicated by a reduction in TMRM fluorescence intensity. The cisplatin-induced reduction in TMRM fluorescence intensity was not associated with changes in neuronal mitochondrial content as determined by labeling mitochondria with the membrane potential-independent dye Mitotracker (Figure 2B). Together these findings indicate that cisplatin reduces neuronal mitochondrial membrane potential without leading to an actual loss of mitochondria under the conditions tested. Co-culture with astrocytes restored the mitochondrial membrane potential of neurons cultured with cisplatin (Figure 2A). Confocal microscopy confirmed that cisplatin reduces TMRM staining in somata and dendrites of neurons and that co-culture with astrocytes restored TMRM staining (Figure 2C). The increase in TMRM fluorescence intensity of cisplatin-treated neurons in response to astrocytes was associated with an increase in mitotracker fluorescence intensity in neuronal cells (Figure 2B), indicating an increase in neuronal mitochondrial content as a result of co-culture with astrocytes.

**Figure 2:**
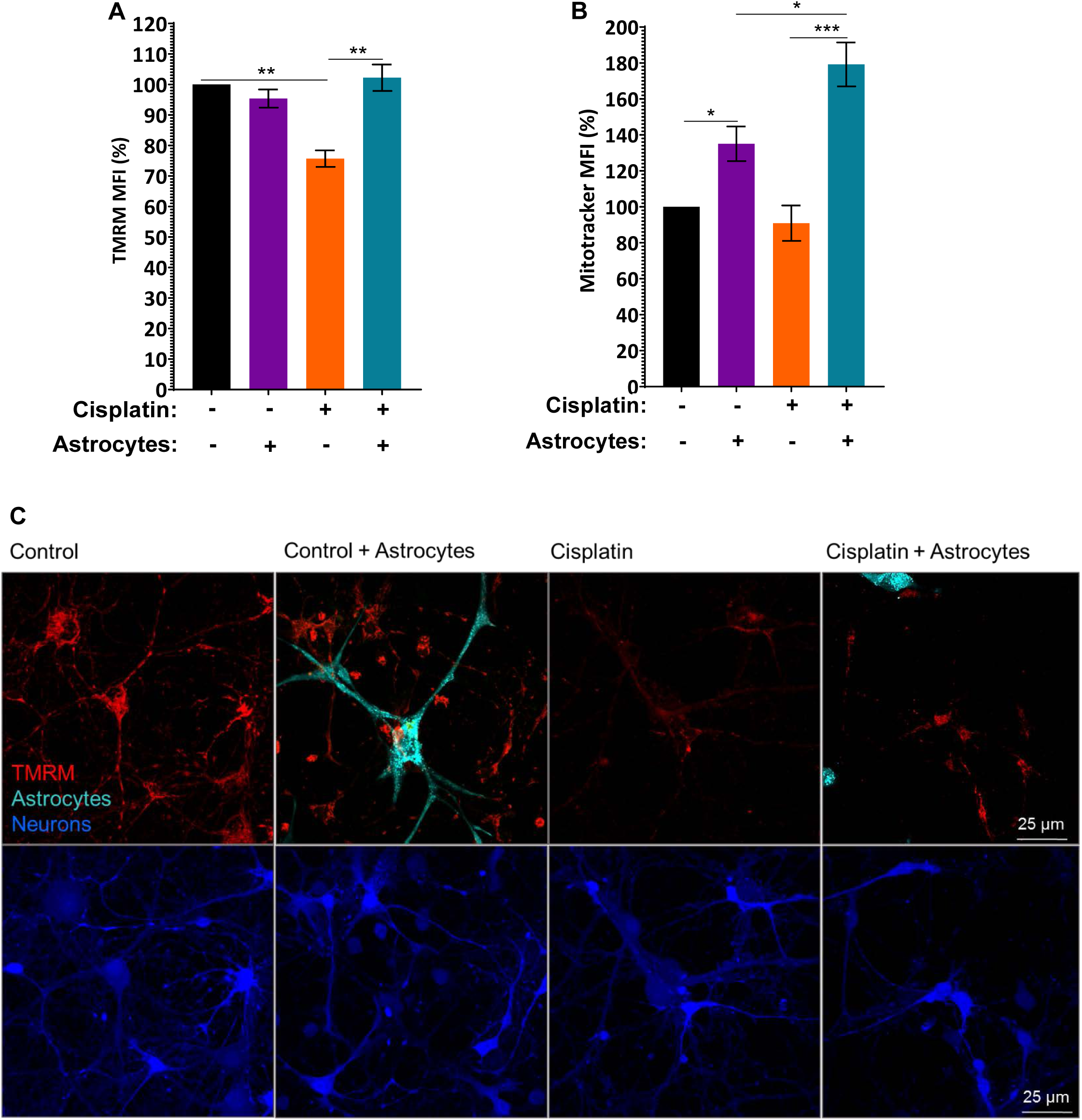
Astrocytes improve mitochondrial membrane potential of neurons damaged by cisplatin. After treating neurons and astrocytes separately with and without cisplatin for 24 h, neurons were labeled with Celltracker Green (CTG) and co-cultured with astrocytes for 17 h. Cells were labeled with TMRM (A) to assess mitochondrial membrane potential or with mitotracker (B) to quantify mitochondrial content by FACS analysis. Fluorescence intensity was normalized to fluorescence intensity in control CTG-positive neurons for each experiment and data represents the mean ±SEM of 3 independent experiments performed in duplicate. Two-way ANOVA TMRM: cisplatin x astrocyte interaction: \**p*< *0*.*05*; Tukey’s post-hoc test: \*\**p*<*0*.*01*. Mitotracker: cisplatin x astrocyte interaction: \**p*< *0*.*05*; Tukey post-test: \**p*<*0*.*05*, \*\*\**p*<*0*.*001*. (C). Representative confocal images of the TMRM signal (red) with neurons labeled with CTB and astrocytes with cell tracker deep red (Teal). Scale bar: 25µm.

### Astrocytes transfer mitochondria to neurons damaged by cisplatin

Next, we tested the hypothesis that astrocytes transfer mitochondria to neurons damaged by cisplatin. We labeled astrocyte mitochondria with mCherry coupled to a mitochondrial localization sequence (mito-mCherry). Neurons and astrocytes were treated separately with cisplatin or control medium for 24 h; neurons were labeled with Celltracker Blue, and co-cultured with Celltracker Deep Red-labeled astrocytes for an additional 17 h. Confocal fluorescence analysis revealed that cisplatin-treated neurons contained astrocyte-derived mCherry+ mitochondria. Only a few astrocyte-derived mitochondria were present in untreated control neurons co-cultured with astrocytes (Figure 3A-C). These findings indicate that astrocytes transfer mitochondria to neurons damaged by cisplatin. Quantitative assessment of mitochondrial transfer showed that cisplatin induced an approximately 3-fold increase in the percentage of neurons that had received mitochondria from astrocytes (Figure 3D). 3D reconstruction and orthogonal section analysis confirmed that the mito-mCherry+ mitochondria are localized inside the neurons and were detected in the soma, axons and dendrites of the neuron (Figure 3E-F).

**Figure 3:**
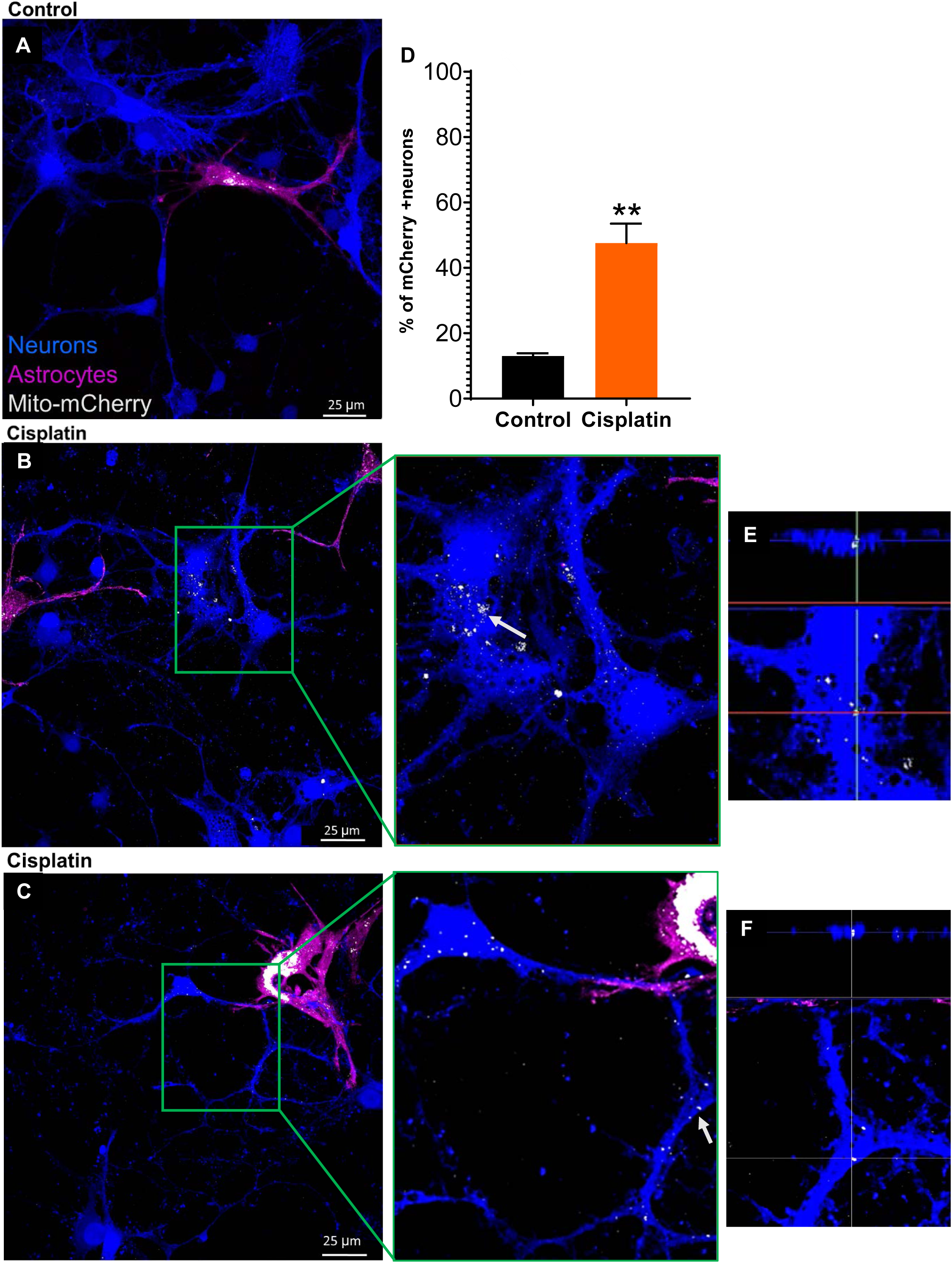
Astrocytes transfer mitochondria to neurons damaged by cisplatin. (A). Representative confocal image of untreated neurons co-cultured for 17 h with astrocytes in which mitochondria were labeled with mCherry. (B-C). Representative confocal images of co-cultures of cisplatin-treated neurons labeled with CTB and cisplatin-treated astrocytes in which mitochondria were labeled with mCherry. Middle panel: larger magnification of the boxed area in B/C. (E-F). 3-D reconstruction and orthogonal slicing showing that the astrocyte-derived mCherry-positive mitochondria (identified by arrows in the middle panel) are present within the neurons. Scale bar: 25µm. (D). Quantification of neurons in co-cultures containing mCherry+ astrocyte-derived mitochondria. Paired Student’s t-test \*\**p*<*0*.*01*. Data represents the mean ±SEM of 3 independent experiments performed in duplicate.

### Role of Miro-1 in mitochondrial transfer from astrocytes to neurons

We next assessed the contribution of Miro-1, a Rho-GTPase mitochondrial adaptor protein, to mitochondrial transfer from astrocytes to damaged neurons. Astrocytes were transfected with short interfering RNA (siRNA) to knock down Miro-1 (astrocytes^Miro-1 siRNA^) and mitochondria in astrocytes were labeled with mito-GFP. Miro-1 siRNA decreased astrocytic Miro-1 by approximately ∼50% (Figure 4 inset). Neurons were treated with cisplatin and co-cultured with astrocytes^Miro-1 siRNA^ or astrocytes^Scr siRNA^ as a control and mitochondrial transfer was quantified. The results in Figure 4 show that knockdown of Miro-1 in astrocytes decreased mitochondrial transfer to neurons in comparison to astrocytes treated with control scrambled siRNA (Figure 4). These findings indicate that Miro-1 plays a key role in the transfer of mitochondria from astrocytes to neurons damaged by cisplatin.

**Figure 4:**
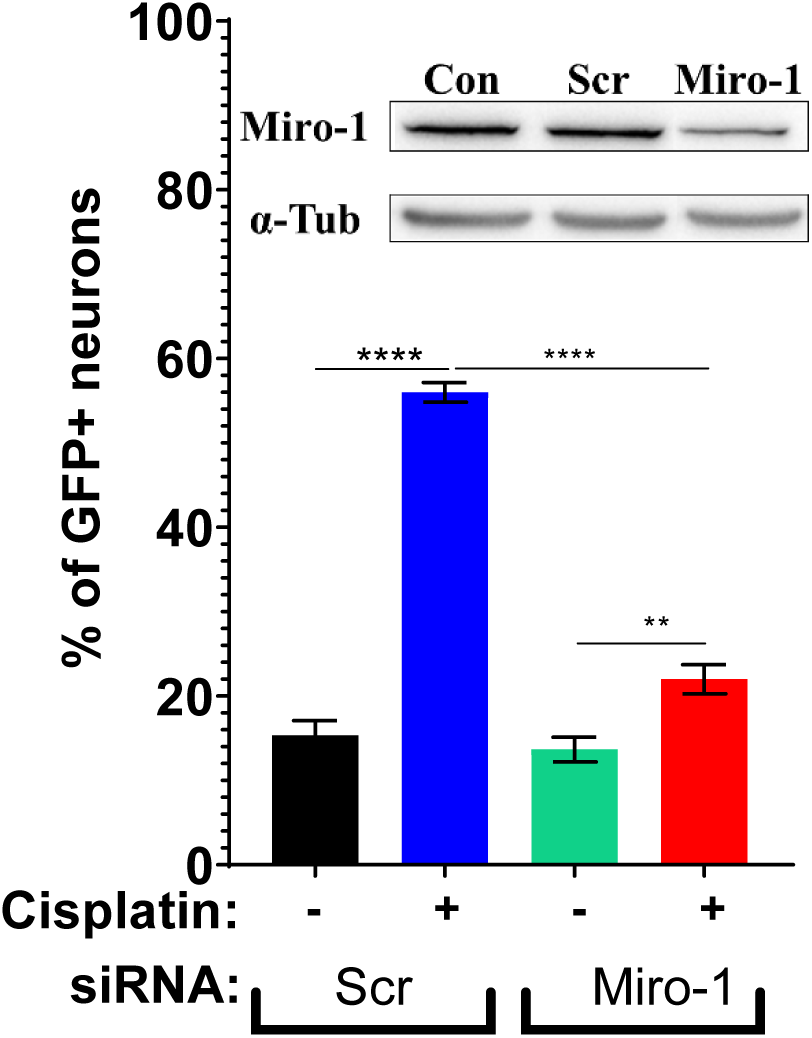
Role of Miro-1 in mitochondrial transfer from astrocytes to neurons. Astrocytes were transfected with mito-GFP and Miro-1 siRNA or control scrambled (Scr) siRNA cultured with or without cisplatin 24 hours and co-cultured for 17 hours with neurons treated with or without cisplatin. Using confocal microscopy the percentage of neurons containing GFP was quantified. Data represents the mean ±SEM of 3 independent experiments performed in duplicate. Inset: Western blot to confirm knockdown of Miro-1 in astrocytes after transfecting with mito-GFP, Scr and Miro-1 siRNA. The percentage of knockdown was normalized to the control and quantified. Two-way ANOVA cisplatin x transfected astrocyte interaction: \*\*\*\**p*<*0*.*0001*, followed by Tukey’s post-hoc test: \*\**p*<*0*.*01*, \*\*\*\**p*<*0*.*0001*.

### Effects of cisplatin and astrocytes on neuronal calcium dynamics

We next asked the question whether cisplatin alters neuronal Ca^2+^ levels and whether co-culture with astrocytes affects cisplatin-induced changes in neuronal calcium dynamics. Resting [Ca^2+^_i_] levels, as measured using the calcium indicator Fura-2, were significantly higher in cisplatin-treated neurons than in control neurons (Figure 5A and B). Co-culture with astrocytes normalized resting [Ca^2+^_i_] levels in cisplatin-treated neurons (Figure 5A and B). The increase in neuronal [Ca^2+^_i_] level in response to 20mM KCl was smaller in cisplatin-treated than in control neurons and this was also normalized by co-culture with astrocytes, indicating that astrocytes normalize calcium dynamics in neurons treated with cisplatin. (Figure 5C and D).

**Figure 5:**
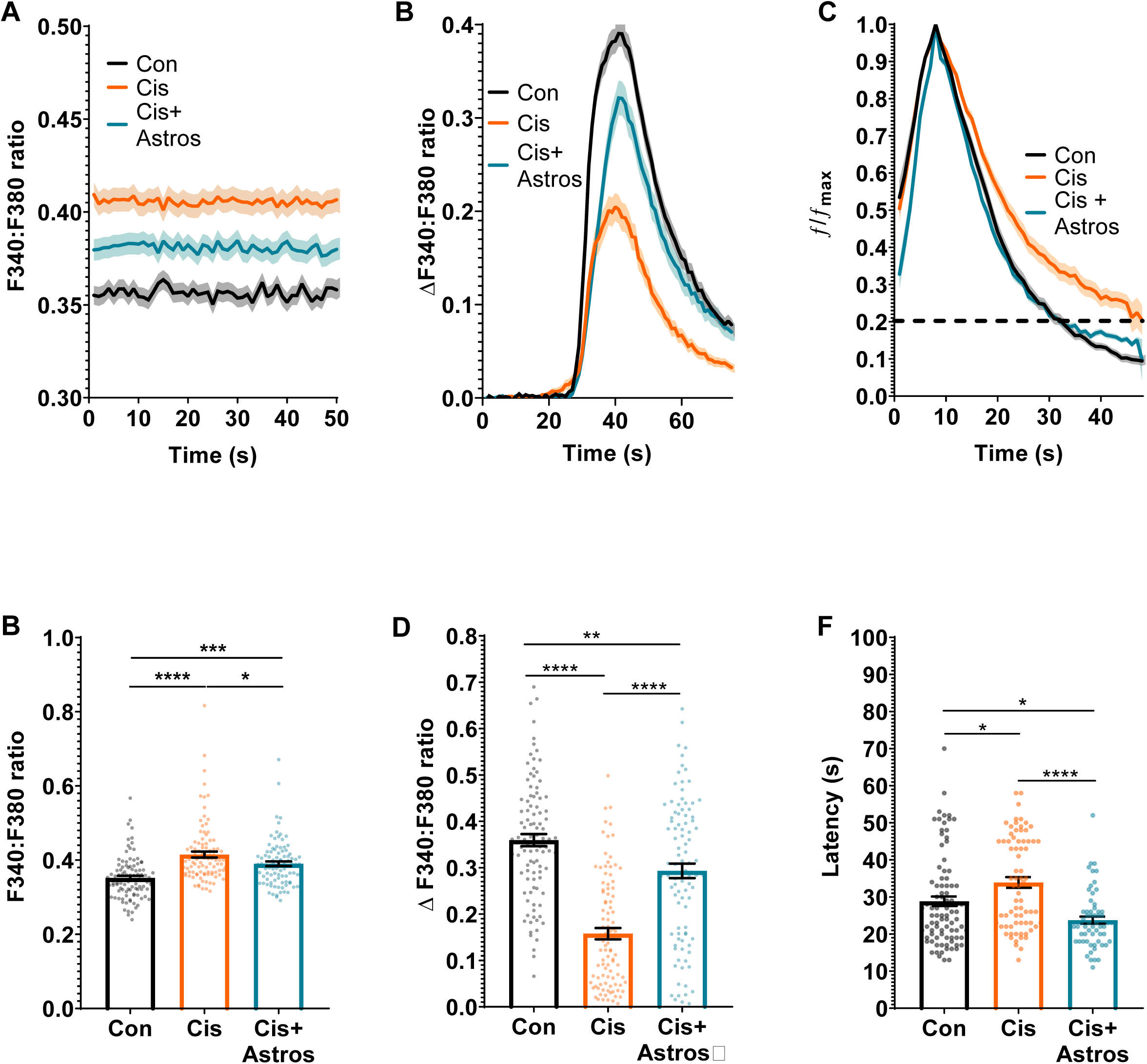
Effects of cisplatin and astrocytes on neuronal calcium dynamics. Neurons were treated with cisplatin followed by co-culture with astrocytes and calcium dynamics were monitored with Fura-2. (A, B). Resting ratio of 340:380 Fura-2 fluorescence intensity (F340:F380) as an indicator of intracellular levels of Ca^2+^ of neurons treated with or without cisplatin followed by co-culture with or without astrocytes. (A). Mean traces and SEM of the F340:F380 ratio of Fura-2 at baseline during first 50 seconds of recording. (B). Mean F340:F380 ratio during 50 seconds of recording for each neuron at rest. Data represents the mean ±SEM of >90 neurons total per group collected in 3 independent experiments. One-way ANOVA, followed by Tukey’s post-hoc test: (\**p*<*0*.*05*, \*\*\**p*<*0*.*001*, \*\*\*\**p*<*0*.*0001*). (C). Mean traces and SEM of the increase in F340:F380 ratio of Fura-2 of neurons responding to stimulation with 20mM KCl. The threshold change in Fura-2 F340:F380 ratio to be considered a response to 20mM KCl was set at an increase of >5 x standard deviations above baseline average. This excluded 9% of neurons in the control group, 23% of neurons in the cisplatin group and 11% of neurons in the cisplatin + astrocyte group. Data represents the mean ±SEM of 3 independent experiments with > 70 neurons for each group. (D). Mean change in F340:F380ratio in response to stimulation of neurons with 20mM KCl. The increase in Fura-2 ratio in response to 20 mM KCl was calculated for each neuron. Data represent the mean ±SEM of n>90 neurons for each group collected in 3 independent experiments. One-way ANOVA, followed by Tukey’s post-hoc test: (\*\**p*<*0*.*01*, \*\*\**p*<*0*.*001*, \*\*\*\**p*<*0*.*0001*). (E) Normalized KCl responses show a delayed decay of Ca^2+^ influx in cisplatin-treated neurons vs. control and cisplatin + astrocytes groups. (F). Quantification of latency to return to 20% of maximal Ca^2+^ (dotted line). Data represents the mean ±SEM of 3 independent experiments with > 70 neurons for each group. One-way ANOVA, followed by Tukey’s post-hoc test: (\**p*<*0*.*05*, \*\*\**p*<*0*.*0001*).

Treatment of neurons with cisplatin increased the time to reach 80% clearance of [Ca^2+^_i_] after exposure to 20 mM KCl (Figure 5E and F), even though maximal calcium levels were significantly lower in cisplatin-treated neurons (Figure 5A and B). These findings indicate Ca^2+^ clearance is impaired in cisplatin treated neurons. Co-culture with astrocytes increased the rate of calcium clearance in cisplatin-treated neurons to a level that was even higher than that of control neurons (Figure 5E and F).

### Mitochondrial transfer underlies the beneficial effects of astrocytes on calcium dynamics in damaged neurons

To address the question of whether transfer of mitochondria from astrocytes to neurons contributes to the improved neuronal Ca^2+^ dynamics, we compared calcium responses to 20 mM KCl of neurons that did or did not contain mCherry+ mitochondria on the same coverslip. As shown in Figure 6A neurons that contained astrocyte-derived mitochondria showed a larger increase in 20mM KCl evoked [Ca^2+^_i_] in comparison to neurons in the same culture that did not receive astrocytic mitochondria (Figure 6A). This finding indicates that transfer of mitochondria from astrocytes to neurons plays a substantial role in restoring the KCl evoked [Ca^2+^_i_] increase in neurons damaged by cisplatin.

**Figure 6:**
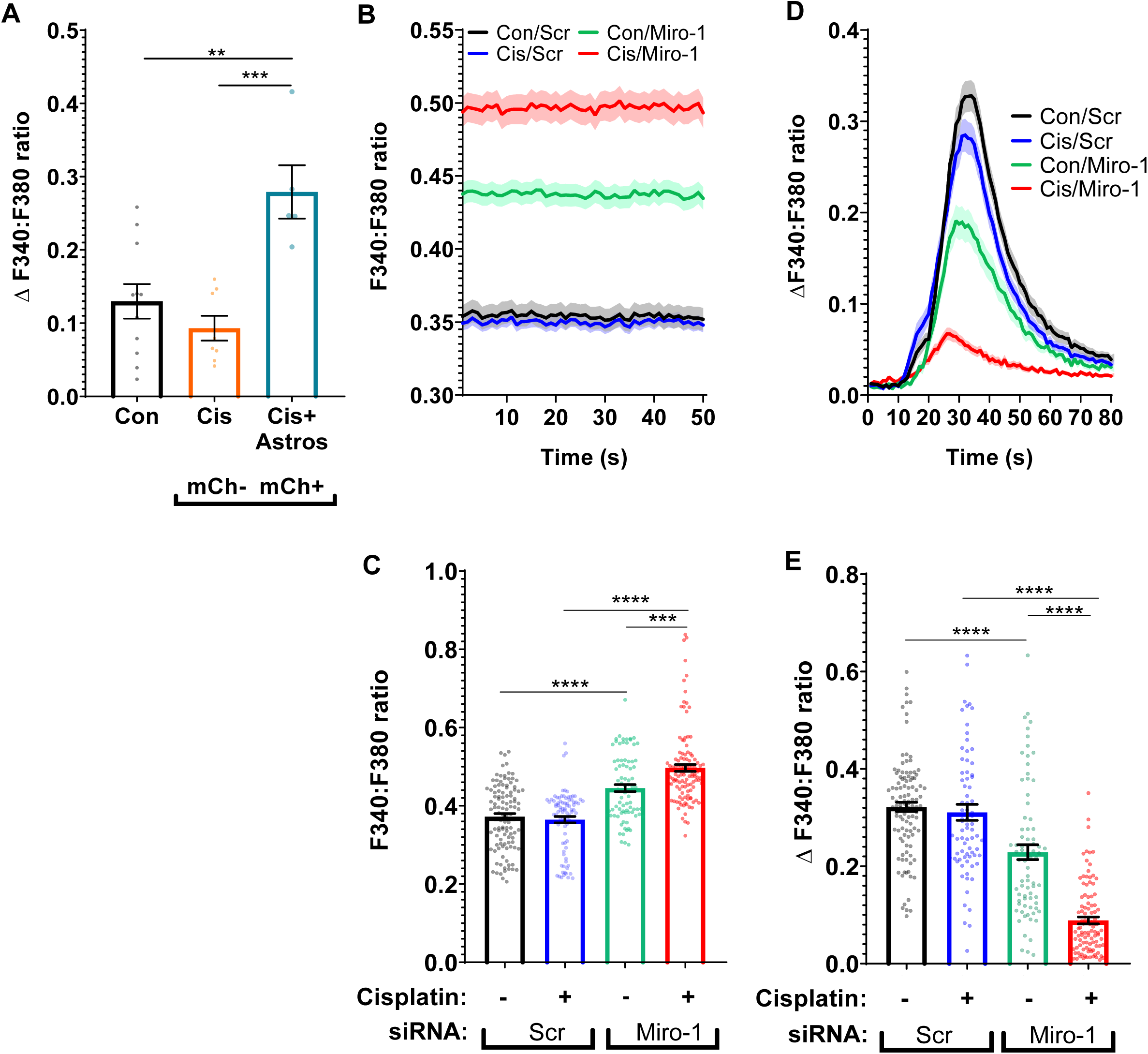
Mitochondrial transfer underlies the beneficial effects of astrocytes on calcium dynamics in neurons damaged by cisplatin. (A). Neurons were treated with and without cisplatin for 24h and co-cultured with astrocytes transfected with mito-mCherry (labeled mCh) for 17h. The calcium response to 20 mM KCl was compared in cisplatin-treated neurons cultured without astrocytes (black bar), or with astrocytes separated into mCherry negative neurons (orange bar; do not contain astrocytic mitochondria) and mCherry positive neurons (blue bar; contain astrocyte-derived mitochondria). The mean increase in the F340:F380 ratio of Fura-2 fluorescence in response to 20 mM KCl was calculated as in figure 5. Data represents the mean ±SEM of 3 independent experiments with n=24 neurons in total. One-way ANOVA, followed by Tukey’s post-hoc test: (**p<0.01, ***p<0.0001). (B). Resting F340:F380 ratio of Fura-2 of neurons treated with cisplatin followed by co-culture with astrocytes transfected with mito-GFP and scr siRNA or Miro-1 siRNA. (C). Mean F340:F380 ratio was calculated for each neuron for first 50 seconds and data represents the mean ±SEM of 3 independent experiments with n>80 neurons for each group. Two-way ANOVA: cisplatin x transfected astrocyte interaction: \*\*\**p*<*0*.*001* followed by Tukey’s post-hoc test: (\*\*\**p*<*0*.*001*, \*\*\*\**p*<*0*.*0001*). (D). Mean traces and SEM of the F340:F380 ratio in response to 20mM KCl stimulus in neurons treated with and without cisplatin and co-cultured with astrocytes transfected with scrambled (Scr) or Miro-1 siRNA. The mean F340:F380 ratio KCl responses were normalized to baseline. (E).The change in F340:F380 ratio between baseline and maximum peak value from the 20mM KCl stimulus was calculated which correlated with [Ca^2+^]_i_ increase in neurons. Mean increase in F340:F380 ratio ratio was calculated for each neuron for 3 sec before 20mM KCl and subtracted from mean F340:F380 ratio at the maximum response to 20mM KCl. Data represents the mean ±SEM of 3 independent experiments with n>80 neurons for each group. Two-way ANOVA: cisplatin x transfected astrocyte interaction: (\*\*\*\**p*<*0*.*0001*), followed by Tukey’s post-hoc test (\*\*\*\**p*<*0*.*0001*).

To further address the contribution of mitochondrial transfer to the normalization of calcium dynamics in neurons in response to co-culture with astrocytes, we assessed the effect of astrocytic Miro-1 knockdown. The results in Figure 6B and C show that basal calcium levels were lower in cisplatin-damaged neurons co-cultured with control astrocytes^Scr siRNA^ as compared to astrocytes^Miro-1 siRNA^. This indicates that knockdown of Miro-1 in astrocytes prevented the normalization of baseline calcium levels in cisplatin-treated neurons (Figure 6B and C). Similarly, co-culture of cisplatin-treated neurons with astrocytes^Miro-1 siRNA^ failed to normalize the 20mM KCl evoked [Ca^2+^_i_] that was observed in the presence of control astrocytes^Scr siRNA^ (Figure 6D and E). This indicates that the restorative effects of astrocytes on calcium dynamics in cisplatin-treated neurons are abrogated by impairing mitochondrial transfer from astrocytes.

## Discussion

Our *in vitro* findings suggest that astrocytic mitochondrial transfer to neurons may represent an endogenous repair mechanism to counteract the neurotoxic effects of cisplatin treatment. Our evidence indicates that cisplatin reduces neuronal survival and decreases mitochondrial membrane potential in the surviving neurons. We also show that co-culture of cisplatin-treated neurons with astrocytes results in mitochondrial transfer from astrocytes to neurons and this is associated with normalization of survival and mitochondrial membrane potential. Cisplatin altered neuronal Ca^2+^_i_ levels in cortical neurons damaged by cisplatin, and addition of astrocytes normalized neuronal Ca^2+^_i_. Calcium levels were specifically restored in those cisplatin-treated neurons that had received astrocytic mitochondria. Moreover, we show that the Rho-GTPase Miro-1 is essential for the transfer of mitochondria from astrocytes to neurons. SiRNA-mediated knockdown of astrocytic Miro-1 prevented transfer of mitochondria from astrocytes to damaged neurons and prevented the restoration of calcium dynamics in neurons damaged by cisplatin. Collectively, our data support the hypothesis that astrocytes counteract the neurotoxic effects of cisplatin by transfering mitochondria to neurons damaged by cisplatin via a Miro-1-dependent pathway.

The effects of astrocytic mitochondrial transfer have been previously shown in disease models of Alexander’s disease and ischemic stroke. Specifically, a prior study using an ischemic stroke model in rodents demonstrated that astrocyte-derived conditioned medium improved the survival of neurons that were damaged by oxygen-glucose deprivation. In our *in vitro* system, we investigated whether astrocytes could also have a restorative effect on neuronal damage as a result of cisplatin treatment. To that end we pre-incubated neurons with cisplatin which caused a significant decrease in neuronal survival and a reduction in the mitochondrial membrane potential in the surviving neurons, and subsequently co-cultured the surviving neurons with astrocytes. Our data show that astrocytes can repair already existing neuronal damage as a result of cisplatin because co-culture with astrocytes restored neuronal mitochondrial membrane potential and protected against further neuronal death.

A structural transport mechanism that has been shown to be involved in intercellular mitochondrial transfer are tunneling nanotubes (TNTs). TNTs are thin non-adherent actin-rich membranous structures with diameters between 50-1500 nm that can span long distances of several hundred nm (Vignais *et al*., 2017). These TNT form direct connections between cells to transport cellular components including cytoplasm, ions, lipid droplets, viral and bacterial pathogens, genetic material and organelles like lysosomes, and last but not least, mitochondria (Li *et al*., 2017; Vignais *et al*., 2017). Multiple authors have described that the mitochondrial Rho-GTPase-1 protein (Miro-1) is a crucial player in intercellular mitochondrial transfer via TNTs. Miro-1 is an outer mitochondrial membrane protein, that binds to Milton, a kinesin/dynein adaptor protein and this promotes mitochondrial motility. We have now expanded this knowledge by showing that decreasing Miro-1 expression in astrocytes decreased the transfer of mitochondria from astrocytes to damaged neurons. Apparently, astrocytic Miro-1 is required for transfer of astrocyte mitochondria to neurons. This could imply that the formation of TNTs for intercellular transport is executed by astrocytes rather than neurons. Jiang *et al*. (2016) have described that the cyclic ADP ribose hydrolase CD-38 plays an important role in astrocytic mitochondrial transfer. Inhibition of CD-38 with apigenin significantly reduced astrocytic mitochondrial transfer and it has been suggested that CD-38 may be involved in TNT formation (Marlein *et al*., 2019). Furthermore, in an *in vitro* myeloma cancer model, myeloma cells were shown to receive mitochondria from non-malignant bone marrow stromal cells through CD-38-dependent tumor-derived formation of TNTs. However, we have to keep in mind that this phenomenon could be specific for tumor cells.

Miro-1 modulates mitochondrial shape in response to cytosolic Ca^2+^ stress, a phenomenon which is distinct from fission and fusion (Nemani *et al*., 2018). Additionally, Stephen et al. (2015), have shown that Miro-1 positions mitochondria in areas within the astrocytic processes that are near neuronal synaptic activity where high energy and Ca^2+^ modulation is necessary. With this in mind it could be possible that astrocytic Miro-1 could be important for positioning mitochondria for transfer in response to neuronal Ca^2+^ changes due to cisplatin damage. It is very well possible that additional activities of astrocytes involved in mitochondrial/cellular health could be of importance as well. However, we specifically observed restoration of [Ca^2+^]_i_ levels in those neurons that actually had received astrocytic mitochondria which implies that mitochondrial transfer plays a crucial role in the restorative effect of astrocytes on neuronal [Ca^2+^_i_] levels and health. Neuronal Ca^2+^ levels are tightly regulated and are critical for many processes in neurons including neurotransmission, depolarization, and synaptic activities. Mitochondria play an important role in controlling Ca^2+^ levels by taking up, buffering and releasing cytosolic Ca^2+^. Cisplatin can negatively alter Ca^2+^ levels such as has been observed in the dorsal root ganglia (Leo *et al*., 2017). Our findings expand this knowledge by showing that cisplatin leads to dysfunctional cortical neuronal Ca^2+^ levels. Abnormalities in resting and KCl-evoked Ca^2+^ increases and clearance due to treatment of cortical neurons with cisplatin were reversed in the presence of astrocytes. As mitochondria are important for neuronal Ca^2+^ dynamics we suggest that astrocytic mitochondrial transfer is key to normalizing the neuronal mitochondrial network. In support of this hypothesis, reducing mitochondrial transfer with Miro-1 siRNA transfection of astrocytes prevented the restoration of neuronal Ca^2+^ dynamics.

Our cell survival studies showed that astrocytes are much less sensitive to cisplatin than neurons. RNA-seq data (Barres *et*.*al*, 2018), show that astrocytes have a higher concentration of the mitochondrial polymerase gamma (γ) in comparison to neurons which is the sole polymerase involved in mitochondrial replication, mutagenesis and repair of mtDNA. Therefore, we suggest that the releatively high activity of polymerase-γ in astrocytes may lead to efficient repair of cisplatin adducts which could contribute to the observed astrocytic resiliency. In addition, astrocytes have a higher concentration of the copper tansporters ATP7a and ATP7b in comparison to neurons and other glial cells (Barres *et*.*al*, 2018). ATP7a and ATP7b are copper transporters that also promote platinum efflux and thereby may contribute to cellular resistance to cisplatin (Kuo *et al*., 2007; Blair *et al*., 2009; Kilari, Guancial and Kim, 2016).

An important translational question still lingers: if astrocytic mitochondrial transfer occurs in the brain after cisplatin treatment, why do patients undergoing chemotherapy still experience neurotoxicity leading to chemotherapy-induced cognitive impairment ? One argument could be that the endogenous restorative capacity of astrocytes is no longer sufficient when patients are treated for a long time, which is common for chemotherapy with cisplatin. Indeed, the risk of developing chemobrain increases with duration of treatment. Moreover, our preliminary data indicate that exposure of mice to a single round of cisplatin treatment does not induce cognitive deficits, whereas two rounds of cisplatin do induce significant decreases in performance in tests of cognitive function (Zhou et al., 2016, Chiu et al., 2016). Although astrocytes may still prevent the actual death of (non self-renewing) adult neurons, they may may fail to completely restore mitochondrial health. The endogenous protective activity of transferring healthy mitochondria from astrocytes to damaged neurons may also become less efficient in the aging brain. From the literature it is known that the severity of the behavioral neurotoxic effects are correlated with age (Asher, 2011; Gutmann, 2019). When endogenous protective mechanisms are not sufficient, interventions aimed at restoring mitochondrial health may provide additional help. Indeed, we showed recently that cell therapy with mesenchymal stem cells or a pharmacolgocial intervention with and HDAC6 inhibitor both reverse cisplatin-induced neuronal mitochondrial abnomarlities as well as cognitive impairment in mice(Chiu *et al*., 2017; Ma *et al*., 2018)

In conclusion, we propose that astrocytic mitochondrial transfer is an important endogenous protection mechanism against chemotherapy neurotoxicity. Promoting astrocytic mitochondrial transfer could represent interesting therapeutic targets to prevent or treat the devastating effects of chemotherapy on the brain.

## Acknowledgments

This study was supported by grants RO1 CA208371 (CJH, AK) and R01 CA227064 (AK, CJH) from the National Cancer Institute. This work was also supported by the NIH/NCI under award number P30CA016672 to MD Anderson Cancer Center.

